# A Novel Protein for the Bioremediation of Gadolinium Waste

**DOI:** 10.1101/2023.01.05.522788

**Authors:** Harvey D. Lee, Connor J. Grady, Katie Krell, Cooper Strebeck, Nathan M. Good, N. Cecilia Martinez-Gomez, Assaf A. Gilad

## Abstract

Several hundreds of tons of gadolinium-based contrast agents (GBCAs) are being dumped into the environment every year. Although macrocyclic GBCAs exhibit superior stability compared to their linear counterparts, we have found that the structural integrity of chelates are susceptible to ultraviolet light, regardless of configuration. In this study, we present a synthetic protein termed GLamouR that binds and reports gadolinium in an intensiometric manner. We then explore the extraction of gadolinium from GBCA-spiked artificial urine samples and investigate if the low picomolar concentrations reported in gadolinium-contaminated water sources pose a barrier for bioremediation. Based on promising results, we anticipate GLamouR can be used for detecting and mining REEs beyond gadolinium as well and hope to expand the biological toolbox for such applications.

## Introduction

Many critical decisions in the clinic today rely on Magnetic Resonance Imaging (MRI). Due to its large magnetic moment and a long electronic relaxation time, gadolinium (^64^Gd) is commonly used to enhance the contrast of the MR image[1, 2]. It is estimated that more than 30 million doses of gadolinium-based contrast agents (GBCAs) are administered to patients annually[3], which are then excreted into the environment. Any unused portions of each bottle/syringe are also discarded, totaling up to several hundreds of tons of GBCAs being emptied into our wastewaters every year. The accumulation of gadolinium in our environment is being addressed with increasing urgency since the second decade of the 21^st^ century; recent studies show anthropogenic gadolinium to be lurking in our wastewaters, crops, tap water, and even marine wildlife[4, 5].

One side effect of administering gadolinium containing products *in-vivo* would be the accumulation inside our bodies. It has been reported that repeated administration of GBCAs leads to gadolinium retention inside the skin, bones, brain, and muscle[6]. The species of which the gadolinium exists (chelated, non-chelated, bound to other compounds such as phosphates, etc.) has yet to be proven with hard evidence. However, should gadolinium break free from their chelates, it would be possible for non-chelated, free gadolinium to bind to receptors which should otherwise be binding calcium[7]. This would hinder any biological functions requiring calcium, such as muscle contraction or neuronal activity – typically found in symptoms associated with Nephrogenic Systemic Fibrosis and Gadolinium Deposition Disease[8-10]. Accumulation of gadolinium in the environment has now been going on for more than 30 years since the introduction of gadopentetate in 1988, with increasing GBCA administration every year.

The current state-of-art for disposing radioactive waste is to collect and store for 10 half-lives, or when activity has decayed to less than 0.1%[11]. Pharmaceutical waste is treated and incinerated at legally registered, licensed facilities[12]. GBCAs, on the other hand, do not have any regulations in terms of proper disposal – the rational being that gadolinium is safe to use when properly chelated. Many GBCAs have been FDA approved to use in the clinic for their safety during bodily circulation and the unparalleled contrast they provide for better healthcare. Although gadolinium retention inside the body has recently come to light, the benefits of GBCAs still undoubtedly outweigh the possible side effects for repeated administration in critical patients[13]. However, the long-term effects of GBCAs being disposed into the environment (and, ultimately, the human body, which consumes products of the environment) have not been fully considered.

As a potential solution to mitigate Rare Earth Element (REE) pollution, several methylotrophic bacteria have been shown to uptake and store light lanthanides and are being proposed as a potential tool for bioremediation and biomining[14, 15]. Furthermore, we have recently developed a mutant of *Methylobacterium*[16]/*Methylorubrum extorquens* AM1 that has acquired the ability of hyperaccumulating heavy lanthanides such as gadolinium*[17]*. The methylotrophic bacteria express a unique protein with the ability to bind lanthanides with picomolar affinity, termed lanmodulin[18, 19]. Lanmodulin binds lanthanides via EF-hand motifs that are 12 amino acids long[20]. Due to the similarity of these lanthanide-binding motifs to calcium-binding motifs, we used the backbone of the well-established genetically encoded calcium indicator GCaMP[21] to construct a new synthetic protein termed Green Lanmodulin-based Reporter (GLamouR), which reports gadolinium binding through green fluorescence.

## Results

### Rapid detection and quantification of gadolinium

Gadolinium detection capabilities of the GLamouR were assessed with fluorescence spectroscopy. Upon binding gadolinium, the protein increases its green fluorescence (Figure 1a). Ten readings were measured and averaged out for each datapoint every ten seconds, and the reaction was found to be within this window of time, suggesting that binding occurs almost instantly. As seen in Figure 1b, the fluorescence increases immediately after injection and does not increase as more time is given, with slight photobleaching resulting from repeated reads. TRIS buffer and calcium were also tested as negative controls, showing a decrease due to the lowered concentration of fluorescent protein upon administration of the negative controls followed by slight photobleaching over consecutive reads. With increasing concentrations of gadolinium in separate wells, we observed the protein’s saturation kinetics. Figure 1c, shows that the GLamouR has a linear response up to 50 μM, suggesting its capability of quantifying gadolinium in the micromolar range. Furthermore, GLamouR is reliably detectable at 10 nM (Figure S1a) and is sensitive enough for 200 nM concentrations of gadolinium (Figure 1d). GLamouR was also tested for its thermal/temporal stability by leaving a 20 μM solution of GLamouR at room temperature for a period of one week. Although there was a slight decline in performance increase over time, GLamouR was able to maintain functionality for the entirety of the week (Figure S1b).

**Figure 1.**
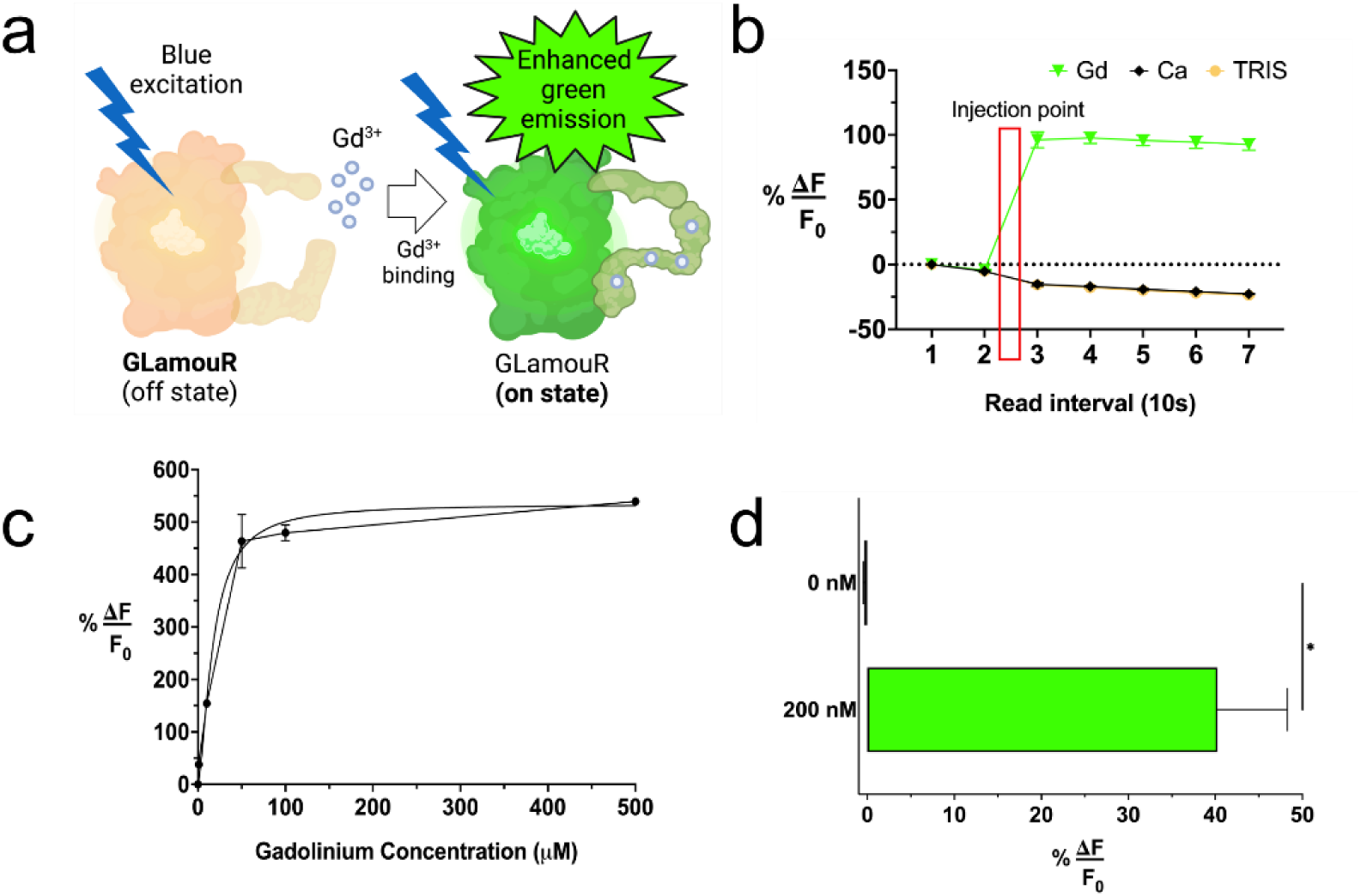
Fluorescence properties of GLamouR. (a) Schematic representation of GLamouR-Gd^3+^ binding. GLamouR in its “on” state will possess enhanced green emission upon excitation with blue light. (b) Fluorescence kinetics of GLamouR with gadolinium (Gd), calcium (Ca), and TRIS buffer injection, which were administered after the second datapoint (ten reads per datapoint). (c) Fluorescence saturation curve for GLamouR with gadolinium shows a linear response up to 50 μM. (d) 200 nM concentrations of gadolinium were detectable with GLamouR with a 40% increase in fluorescence upon injection. Statistical significance was calculated by an unpaired t-test with Welch’s correction, p value=0.0128.

### Verification of gadolinium binding via membrane filtration and MRI

Membrane filtration and MRI were used to verify that GLamouR was in fact capturing gadolinium and not just reporting its presence in an indirect manner. After binding GLamouR with gadolinium, the solution was dialyzed to remove excess gadolinium and subsequently filtered with washed membranes to isolate the protein (Figure S2). The retentate would therefore hold GLamouR-Gd conjugates while the filtrate would be the surrounding buffer. T1 and T2 relaxation times are shown in Figures 2a and 2b, with their corresponding T1 and T2 maps (Figures 2c, 2d). Figures 2a-d show the filtrate containing insignificant levels of gadolinium after dialysis, whereas the protein was able to hold on to gadolinium at high levels of centrifugal force and possess MRI contrast. By plotting relaxation rates as a function of gadolinium concentration, we obtained the r1 and r2 relaxivities of GLamouR-Gd conjugates, which were 6.0 mM^-1^s^-1^ and 41.85 mM^-1^s^-1^ respectively. These numbers represent the enhanced relaxivity of gadolinium upon binding GLamouR (SM, Table1) [22-26], which is likely to be resultant from the tumbling time shifting closer to the Larmor frequency as a conjugate, when compared to free gadolinium. To obtain higher concentrations of GLamouR-Gd conjugates, GLamouR was lyophilized, reconstituted at a lower volume with DDIW and dialyzed with fresh TRIS buffer at 25 mM to match the salt concentrations before initiating gadolinium binding. Compatibility with centrifugal force, lyophilization, and long dialysis times demonstrate the durability and stability of GLamouR. Thermal properties were also assessed with a thermal shift assay suggesting tighter folding at temperatures closer to 45°C, but minimal variance in performance was found due to temperature (Figure S3).

**Figure 2.**
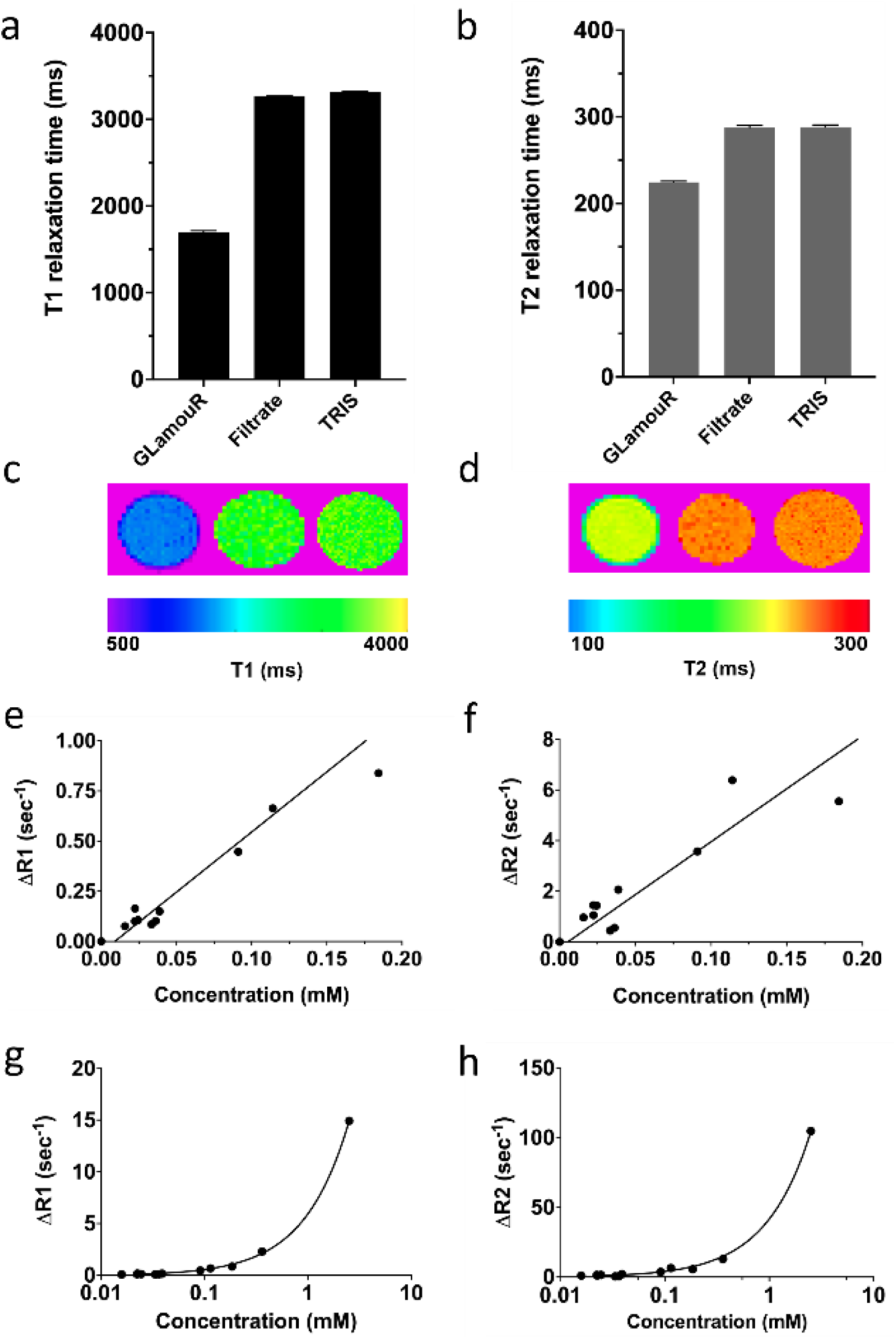
MRI data confirm protein-Gd binding. GLamouR was bound with gadolinium and dialyzed thoroughly prior to separation of filtrate (surrounding buffer) and retentate (GLamouR-Gd conjugates). T1 (a) and T2 (b) relaxation times were compared against TRIS buffer by averaging the ROIs of their corresponding T1 (a) and T2 (b) maps. r1 and r2 relaxivity (per-gadolinium) were derived from thirteen dilutions across six different batches of GLamouR. Linear regression for R1 and R2 datapoints as a function of gadolinium concentration are shown in linear (e, f) and logarithmic scale (g, h), with p < 0.00001 and r^2^ = 0.999

### GBCAs used in the clinic today are susceptible to breakdown from UV irradiation regardless of configuration

Macrocyclic chelates have been proven to be more stable than linear chelates, and thus considered safer to use[27]. Although GBCAs are used immediately upon opening in clinical practice, it is not uncommon for researchers to keep bottles of GBCAs for prolonged periods of time for *in-vitro* studies.

In Figure 3a, various contrast agents that had been open for an unknown period of time were tested for their stability. GLamouR was used to detect unchelated free gadolinium from unstable chelates that had released gadolinium. As expected, macrocyclic agents had less free gadolinium than linear configurations, with Gadopentetate Dimeglumine triggering the highest fluorescence increase (which is currently discontinued for use in the clinic). However, upon 24 hours of UV irradiation, the GBCAs had similar amounts of free gadolinium in solution regardless of configuration (with the exception of Gadopentetate Dimeglumine, which was much more susceptible to UV breakdown than other chelates).

**Figure 3.**
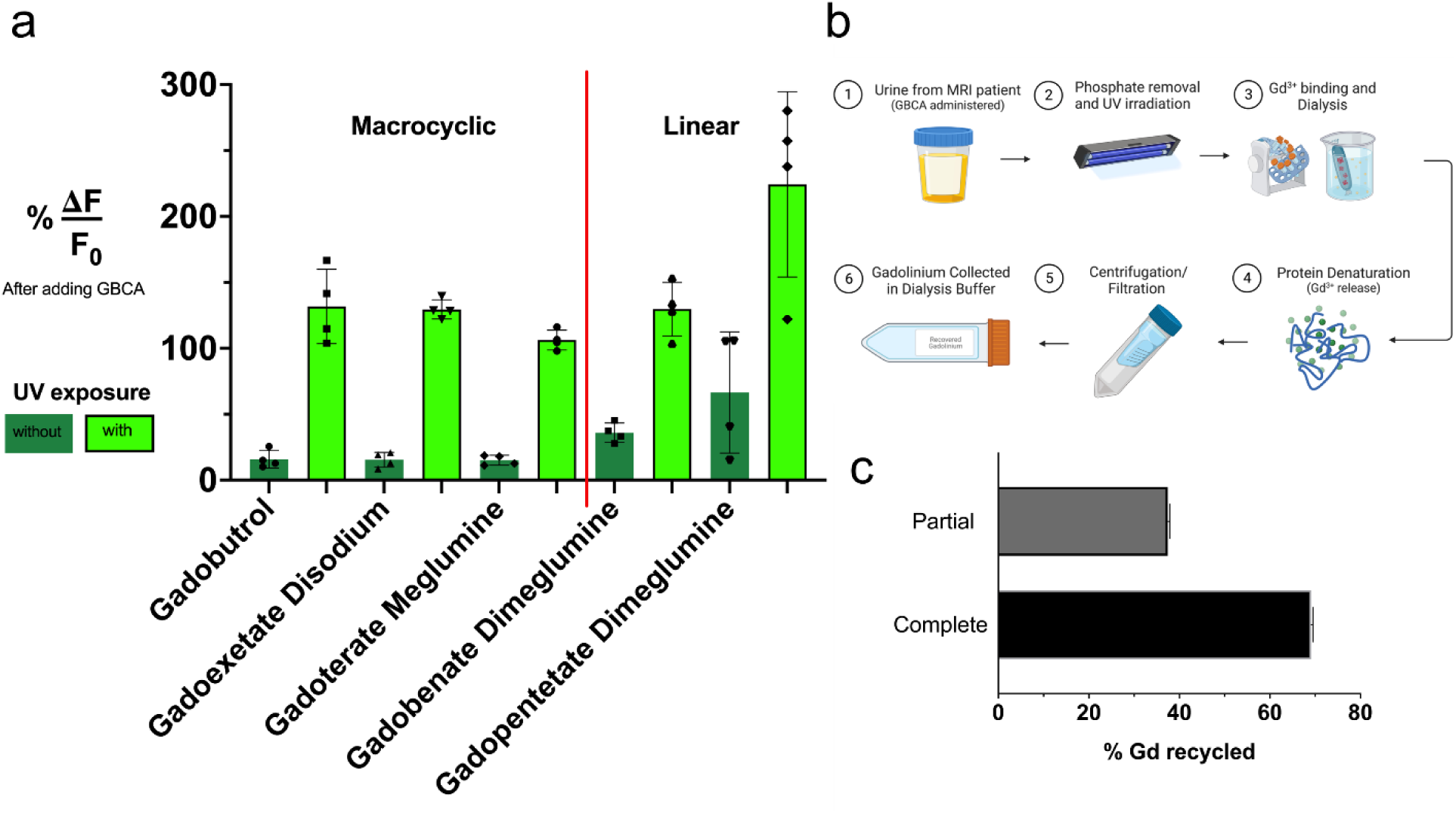
Collection of gadolinium from clinical GBCAs. (a) Clinical GBCAs were tested for unchelated gadolinium ions with GLamouR, before and after UV irradiation. (b) Schematic showing the proposed procedure for collecting gadolinium from MRI patient urine samples, which was substituted for Gadoterate Meglumine-spiked artificial urine in this study. (c) Recycled gadolinium after partial removal and complete removal of phosphates from Gadoterate Meglumine-spiked artificial urine samples. For complete removal, Gadoterate Meglumine was added after passing artificial urine through phosphate removal resin, whereas partial removal was emulated by adding Gadoterate Meglumine before passing through the column with a lower resin ratio.

To emulate urine samples from GBCA administered MRI patients, artificial urine was spiked with the most stable GBCA of the group (Gadoterate Meglumine) to a concentration of 2.25 mM, equivalent to a dose for a 50 kg patient with 90% of GBCA eliminated in 2 L (Figure 3b) for conservative calculation. Samples were then pre-treated by removing phosphates which compete with GLamouR for gadolinium binding, prior to UV irradiation. Once gadolinium had been released from their chelates, GLamouR was introduced to the solution for binding the released ions, which was then subsequently dialyzed to exclusively obtain gadolinium-protein conjugates. After the protein is denatured to release gadolinium, the solution can then be filtered so the final product is gadolinium in TRIS buffer. Figure 3c shows that 37% of total gadolinium was bound to GLamouR with partial removal of phosphates, whereas 69% was bound when they were removed more efficiently, showing that pre-treatment of samples is an important factor for effective extraction.

Based on data represented in Figure 3a, we have found that gadolinium is most likely released from their chelates once GBCAs are excreted into the environment, especially in circumstances involving prolonged UV irradiation from the sun. However, we have also demonstrated that gadolinium can potentially be collected from GBCA-administered MRI patient urine samples before they get excreted into the environment, which can be a preventative measure for such pollution.

### Collecting, detecting, and recycling low concentrations of gadolinium waste with affordable technology

Gadolinium has been reported to exist at the picomolar scale in various polluted bodies of water, even including tap water[4]. To demonstrate the possibility of gadolinium collection from environmentally relevant concentrations, gadolinium was introduced into 1 L of TRIS buffer to a concentration of 200 pM. GLamouR was then used to collect the gadolinium, which was subsequently concentrated down to 100 μL. Final gadolinium concentration was measured via ICP-MS and total number of moles was calculated to be 150.07 picomoles, equivalent to 75% of the initial number of moles of gadolinium. Proper pre-treatment is indeed important to obtain a high percentage of collection, as seen in Figure 3c. However, it is demonstrated in Figure 4a that low concentrations of gadolinium do not pose as a hurdle in collecting or filtering anthropogenic gadolinium from water.

This process involves multiple filtration steps, many plastic consumables, and a centrifuge. We therefore designed a device for collecting and detecting gadolinium which further reduces such labor and need for additional resources. Figure 4b depicts a device setup in which the sample would flow through a pre-treatment chamber for removing competing compounds such as phosphates, which may then also include UV-irradiation to break any pre-existing bonds that could interfere with gadolinium collection. GLamouR column would be placed inside the detection unit, where increase of fluorescence would indicate remaining gadolinium, and saturation of fluorescence would indicate the need for a fresh column. The flow of sample would be continuously repeating until fresh columns do not increase in fluorescence, and remaining waste could then be redirected for disposal. The existing columns could then be further processed to release gadolinium as depicted in Figure 3b, steps 4-6. The advantage of this system would be that extremely low concentrations of gadolinium could be concentrated to higher concentrations and subsequently eluted for isolating gadolinium.

The prototype device was constructed with a budget for less than 500 USD (Figure 4c). The outer shell was 3D-printed, with the bulk of the cost coming from the Arduino microcontroller and Hamamatsu spectrometer. Samples can be pumped in and out of the device by creating a pressure differential inside a bottle containing the sample by utilizing an air pump inflator typically found on sphygmomanometers. Figure 4d shows data collected from the device, where gadolinium concentration was brought up to 100 μM by injecting a 1 mM solution of gadolinium equaling 1/10^th^ of the final volume.

### Identification and collection of REEs beyond gadolinium

Based on Figures 1-4, GLamouR is a novel candidate protein for the detection, quantification, and recycling of gadolinium waste. However, gadolinium is not the sole REE pollution resulting from industrial activities[28], and thus further experiments were carried out to investigate the protein’s responses to different REEs. As seen in Figure 5b, GLamouR was effective in detecting 11 REEs and distinguishing them from calcium. Change in fluorescence was not particularly unique for different REEs, as most of them lingered around the 100% region. A red-shifted GLamouR (GLamouR-rs, Figure 5a) was then created to see if REEs could be identified due to unique REE responses when used in conjunction with green. While the GLamouR-rs has a much lower brightness in both its on and off states than the original green version, it produces a much higher delta across all REEs with the exception of Lanthanum (Figure 5b), where response was recorded to be very minimal (+8.26%). In this case, green fluorescence could be measured to quantify the REE, while the lack of fluorescence increase in red could specify Lanthanum.

**Figure 4.**
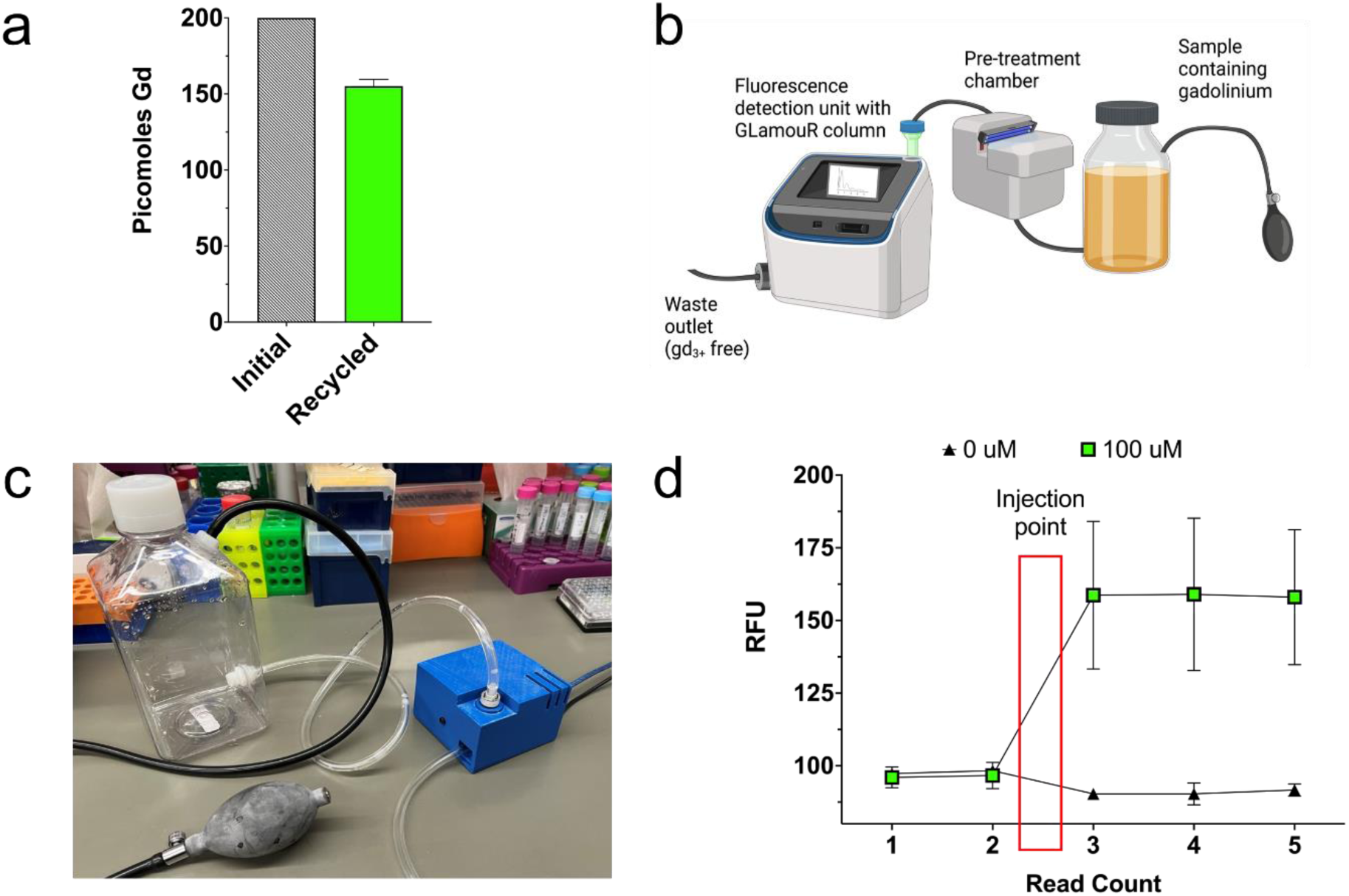
Collection and concentration of picomolar concentrations suggest GLamouR can be used to recover gadolinium from natural water sources. (a) Gadolinium scavenging capabilities at 200 pM (measured with a multimodal plate reader). Gadolinium was gathered and subsequently concentrated down to nanomolar concentration while retaining 75% of total gadolinium mass. (b) Schematic diagram showing the proposed system for mining gadolinium with affordable technology. (c) Photo of the actual prototype built in the lab, to demonstrate the simplicity of the technology and affordable price point. (d) Data produced with the prototype device for gadolinium detection.

**Figure 5.**
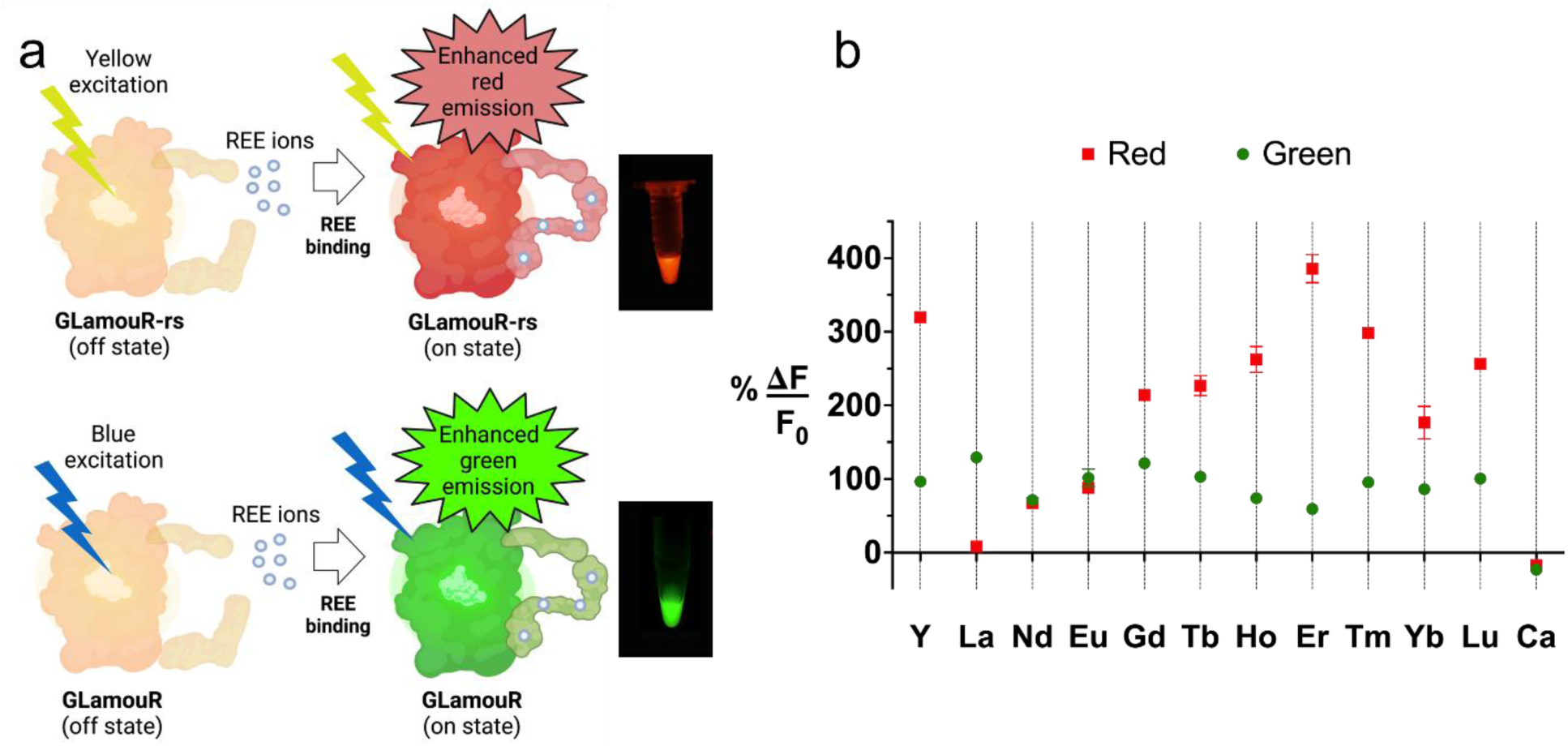
Comparison of GLamouR-rs and GLamouR. (a) Schematic of the red-shifted GLamouR (GLamouR-rs) and original GLamouR binding REEs, with their corresponding fluorescence images to the right. (b) Fluorescence increase of green and red shifted GLamouR upon addition of eleven different REEs show a difference in their preference for certain REEs such as lanthanum (calcium added as negative control).

The creation of GLamouR proteins with unique responses to different REEs suggest the possibility of future GLamouR variants that bind with a stronger affinity or react more efficiently to specific REEs. This would make it feasible to utilize the system depicted in Figure 4c for collecting and isolating specific REEs from environmental samples that contain a mixture of several REEs, as it is often the case in nature. Even with moderate specificities, low concentration mixtures could be accumulated to high enough concentrations for downstream separation to achieve higher yields for cost effectiveness with minimum impurities of undesirable REEs.

In addition to mining and recycling specific REEs, it is hoped that GLamouR variants will allow users to determine REE composition and identify REE-rich sources in a rapid, affordable, and simplistic manner. Although urine samples from GBCA-administered patients have a pre-defined matrix with a guaranteed range of gadolinium concentration, this would not be the case for samples with diverse and unique compositions for every batch, such as electronic waste or samples obtained from the environment.

## Discussion

REEs are an essential resource for modern technology – anything with a screen, lens, glass, lights, magnets, steel alloys, or batteries require the use of REEs, as well as conventional “non-electric” vehicles that run on a form of fossil fuel (which also require a catalytic converter) [29]. Current methods for mining REEs require extensive use of harsh chemicals and intense labor, not to mention low yields and excessive byproducts[30]. Moreover, not only do REEs exist on Earth in a finite amount, but their distribution is alarmingly reliant on REE-rich countries with affordable labor costs and looser regulations on environmental pollution arising from industrial practices[31]. On the bright side, efforts are being made to develop new technologies for mining REEs at a lower cost and for recycling them[32]. The market for recycling REEs was evaluated at 248 million USD in 2021, and it is projected to grow at a rapid pace with a CAGR of 11.2%, reaching 422 million USD by 2026[33].

It would thus be beneficial for nations to develop technology for recycling REEs, preferably at higher yields, cheaper price, and hopefully with environmentally friendlier methods. In addition to utilizing microbes, cutting-edge technology involving the use of peptides and proteins for biomining are being developed by researchers as well [34-40]. Aspects to consider for industrializing the application of such technologies would be the identification of reliable REE sources, performance and reproducibility across various sample types, and cost effectiveness. In this study, we have demonstrated gadolinium recycling from artificial urine at a physiologically accurate GBCA concentration, and collection and concentration of gadolinium from picomolar to nanomolar while retaining roughly 70% of total gadolinium mass for both instances. We have also shown that gadolinium can be detected with a simple setup including a spectrometer, LED light source, and a microcontroller inside a 3d-printed housing for less than 500 USD, making the technology widely accessible and scalable. GLamouR has been evaluated to be a highly stable protein by retaining function for the entirety of a week-long assay at room temperature (SI, Figure 3), while being compatible with lyophilization and high centrifugal force.

Once established on Earth, it is anticipated that GLamouR-like proteins could be used in outer space for extraterrestrial biomining. The near side of the Earth’s moon is reported to possess massive reserves composed of a mixture of Potassium, REEs, and Phosphorus (KREEP), estimated to be 220 million cubic kilometers[41]. A recent study by Cockell et.al., demonstrated extraterrestrial biomining of REEs where microorganisms were tested for their mining capabilities from basaltic rock in various micro-g environments on the international space station[42]. As it is for any type of travel but especially air and space travel, reducing cargo weight is essential for increasing distance per unit fuel. Compatibility with lyophilization to reduce weight during takeoff, simple mining procedures with light and affordable equipment, and instant feedback during REE collection could be desirable characteristics of GLamouR for REE mining in outer space.

The use of biological tools to carry out functions otherwise achieved through chemical, electrical, or mechanical means are often met with higher performance and cheaper production costs, such as the modern development and production of recombinant human insulin in E. coli [43]. Furthermore, biological therapies are the best-selling drugs on the market today [44], and new products that mimic (biosimilars) and even improve upon the precision and efficacy of traditional medications (biobetters) are being developed as well [45]. Such technologies involving biological tools are expected to be tailored for their respective applications in each field, whether it be industrial practices, agriculture, precision medicine, or even unearthing new scientific discoveries.

## Funding

A.A.G acknowledges financial support from the NIH/NINDS: R01-NS098231; R01-NS104306 NIH/NIBIB: R01-EB031008; R01-EB030565; R01-EB031936; P41-EB024495 and NSF 2027113. This work was supported under the MTRAC Program by the State of Michigan 21st Century Jobs Fund received through the Michigan Strategic Fund and administered by the Michigan Economic Development Corporation. The authors wish to thank Dr. Jon Debling for his valuable input.

## Authors contributions

H.D.L and C.J.G were involved in conceptual planning, H.D.L, K.K., C.J.G, and C.S. performed experiments, H.D.L, N.M.G, N.C.M-G, and A.A.G provided methodology, and H.D.L and A.A.G prepared the original manuscript. All authors contributed to the article and have approved the submitted version.

## Competing interests

Authors declare no competing interests.

## Data and materials availability

All data are available in the manuscript or the supplementary materials.

## List of Supplementary Materials

Materials and Methods

Figures S1-S3

Table 1

## Notes

### Competing Interest Statement

The authors have declared no competing interest.

### Summary of Updates

Figure numbers corrected.

