## Supplementary Materials for "A Novel Protein for the Bioremediation of Gadolinium Waste"

**The PDF file includes:**

Materials and Methods
Figures S1-S3
Table 1

### Materials and Methods

Materials

All materials were purchased from Thermo Fisher or its subsidiaries, with the exception of REEs and calcium chloride which were purchased from Sigma-Aldrich, centrifugal filtration units purchased from Merck, and FPLC columns from Cytiva.

Cloning

GLamouR proteins were cloned via Gibson Assembly or TOPO cloning of DNA fragments obtained from PCR of existing constructs (GCaMP6m, rGECO1, Lanmodulin) and custom ordered fragments from IDT. Cloning results were confirmed after transforming the final vector into TOP10 to obtain sufficient DNA for Sanger sequencing (provided by Genewiz) and other downstream applications.

Protein Expression

Proteins were expressed by E. coli (BL21*) that had been transformed with the cloned pET101 vectors containing the GLamouR constructs. Cells were incubated with ampicillin-spiked (100 µg/mL) Magic Media for 24hrs at 30$^{o}$C, shaking at 300-360 RPM. Expression and purification were verified via Western Blot against the V5 tag.

Protein Purification and buffer exchange

Purification was performed via HIS-tag purification with cobalt resin. For small (<50 mL) cultures, 200 uL columns were used, whereas larger volumes (>400 mL) were purified via FPLC (AKTA by Cytiva). Buffer exchange was done with either centrifugal filtration units (3-10kD, 4-15 mL), desalting columns (7kD), or dialysis cassettes (10kD) at least three consecutive times with 25 mM TRIS buffer at pH 7.0. Further purification via size exclusion was performed as necessary, with HiLoad 16/600 Superdex 200pg columns connected to the FPLC system.

Fluorescence Measurements

Fluorescence was measured with the Cytation5 (Biotek) with excitation at 488 nm and emission at 510 nm, with monochromators and/or filters. Wells were prepared with a 10-200 nM concentration of GLamouR (quantified via sequence-specific a205) in TRIS buffer (25 mM, pH7); after the second read, REEs/negative controls were introduced to reach desired concentrations (with ten averages per read).

Magnetic Resonance Imaging

MRI data were acquired using a 7 Tesla 70/30 Biospec (Bruker) equipped with 12-inch gradients and an 86 mm transmit/receive volume coil. T1 maps were generated via Paravision 360, with ten TR experiments ranging from 100 ms-17500 ms with an echo time of 6.89 ms and three averages each. Multiple 1 mm-thick slices were acquired across samples contained in PCR tubes, 0.5 mL tubes, and 1.5 mL microcentrifuge tubes. Samples were prepared by allowing GLamouR (volumes from 400 uL-1 mL, concentrations from 0.5 mg/mL-1 mg/mL) to bind with gadolinium (0.5 mM) for 30 mins at RT, followed by several washing steps, identical to the methods used for buffer exchange following protein purification.

Inductively Coupled Plasma

ICP data were acquired with ICP-OES (Varian 710-ES). Samples were prepared by digestion with nitric acid and final concentrations were calculated based on dilution factors used for each sample.

Relaxivity calculation

Relaxivity ($r_i$) values were calculated with the following equation(*24*) :

$$R_i = R_i^0 + r_i \times [CA]; \qquad i = 1,2$$

where concentration is denoted by [CA], which is the Gadolinium concentration measured by ICP. $R_i$ is the relaxation rate of the solution in presence of the contrast agent (=1/$T_i$). $R_i^0$ is the relaxation rate of the solution without the contrast agent.

Statistical analysis and software

Prism (GraphPad) was used for generating graphs and performing statistical analysis. SnapGene was used for designing vectors containing new proteins and generating their corresponding maps, and BioRender was used for creating illustrations.

**Figure S1.**

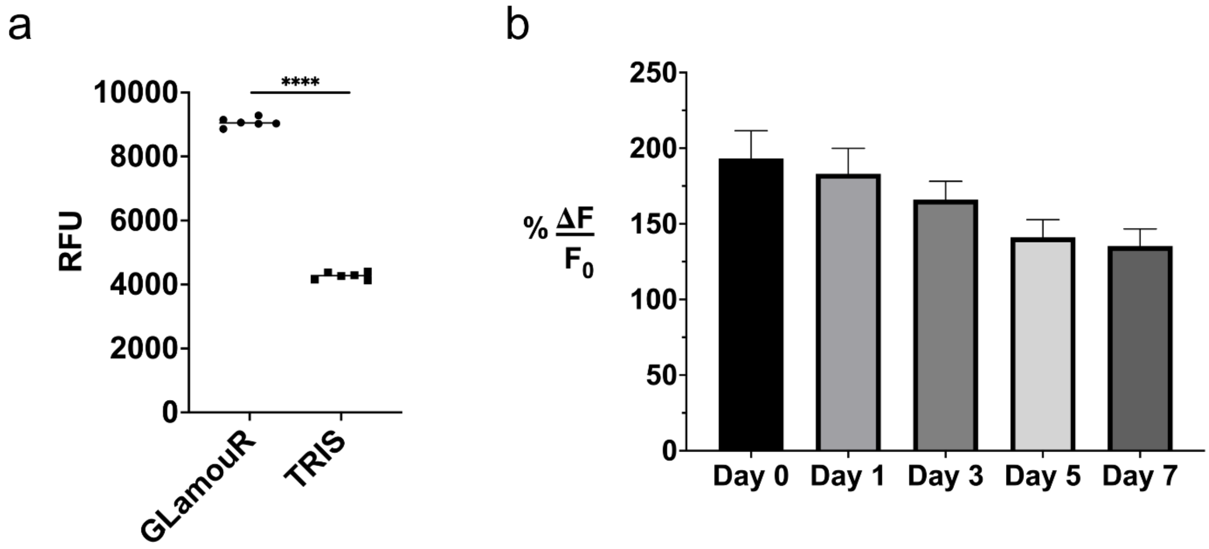


**Baseline readings and thermal/temporal stability of GLamouR.** (a) GLamouR can be reliably detected at 10 nM (n=6, $p < 0.0001$ with Welch's correction) in TRIS buffer and used as baseline readings for assays. (b) GLamouR solutions at 1mg/mL were left at room temperature for up to 7 days and frozen accordingly before thawing for the assay. 2 µL of GLamouR mixed with 243 µL of TRIS buffer were used per well, along with 5µL injections of 5mM gadolinium (n=5).

**Figure S2.**

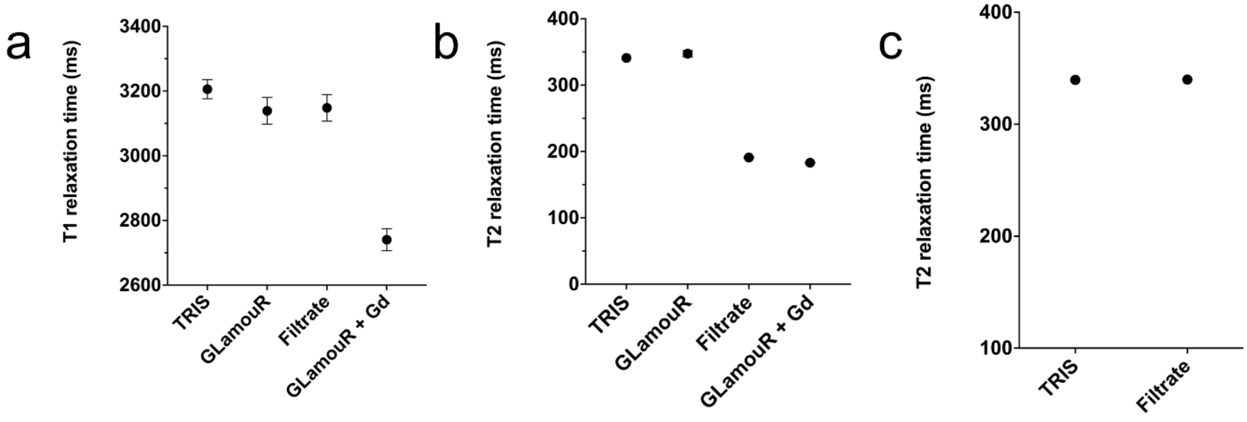


**Washing membranes for filtration and MR Imaging.** The effect of washing membranes before filtration can be seen with MRI. (a) T1 relaxation time does not seem to be lower for the filtrate sample. (b) On the other hand, T2 relaxation time is significantly decreased. (c) Upon washing the membrane before filtration, the filtrate T2 is returned to its normal values. It was found that glycerin on the membranes (for moist storage) were contributing to T2, which could be easily washed off by running DDIW through the membrane.

**Figure S3.**

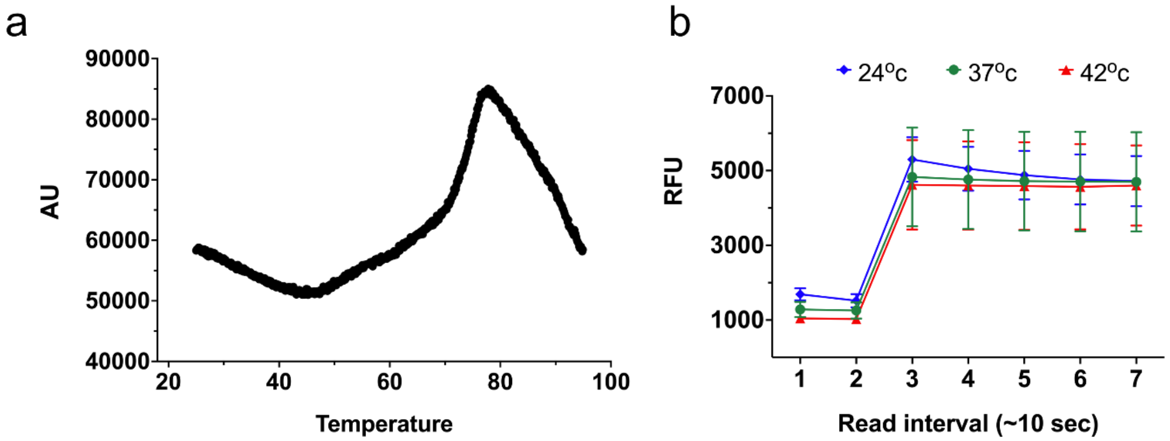


**Thermal properties of GLamouR.** (a) Thermal shift assay was conducted for GLamouR. SYPRO orange dye binds to hydrophobic regions of protein which are increasingly exposed upon denaturation, until aggregation takes place. (b) Performance of GLamouR in different temperatures show minimal variance across tested conditions.

**Table 1.**

| Compound | Buffer | Temperature | Field Strength | r1 (L/mmol-s) | Reference |
|---|---|---|---|---|---|
| GLamouR | TRIS 25mM, pH7 | RT | 7 T | 6.0 | * |
| GdCl3*6H20 | TRIS 25mM, pH7 | RT | 7 T | 4.6 | * |
| Gadoxetate Disodium | Water | 37 °C | 4.7 T | 4.9 | (*26*) |
| | Water | 37 °C | 3 T | 4.3 | (*26*) |
| | TRIS 50mM, pH8 | RT | 7 T | 4.9 | * |
| Gadobutrol | Water | 37 °C | 4.7 T | 3.2 | (*26*) |
| | Water | 37 °C | 3 T | 3.2 | (*26*) |
| | Human plasma | 37 °C | 7 T | 3.8 | (*25*) |
| | Human plasma | 37 °C | 7 T | 4.7 | (*23*) |
| | Human blood | 37 °C | 7 T | 4.2 | (*22*) |
| | TRIS 50mM, pH8 | RT | 7 T | 3.7 | * |
| Gadobenate Dimeglumine | Water | 37 °C | 4.7 T | 4.0 | (*26*) |
| | Water | 37 °C | 3 T | 4.0 | (*26*) |
| | TRIS 50mM, pH8 | 37 °C | 7 T | 3.6 | * |
| Gadoterate Meglumine | Water | 37 °C | 4.7 T | 2.8 | (*26*) |
| | Water | 37 °C | 3 T | 2.8 | (*26*) |
| | Human plasma | 37 °C | 7 T | 2.8 | (*25*) |
| | Human plasma | 37 °C | 7 T | 3.2 | (*23*) |
| | Human blood | 37 °C | 7 T | 2.8 | (*22*) |
| | TRIS 50mM, pH 8 | RT | 7 T | 3.5 | * |

**Relaxivity values for GLamouR-Gd conjugates compared to gadolinium chloride hexahydrate and other chelators.** Asterisks (*) denote values calculated with data collected from this study.
